# Face-selective responses correlate with deep networks that learn from environment feedback

**DOI:** 10.64898/2026.02.25.703652

**Authors:** Mo Zhou, Emily Schwartz, Arish Alreja, R. Mark Richardson, Avniel Ghuman, Stefano Anzellotti

## Abstract

Deep neural networks have shown high accuracy in modeling neural responses in the visual system, but most models rely on supervised learning, which requires training on ground-truth labels that are typically unavailable in real-world settings. While unsupervised models can address this limitation, they miss another key aspect: visual representations are shaped by feedback from the environment. We introduce a reinforcement learning (RL) model of face perception that incorporates both input stimuli and feedback from the environment. Inspired by human interactions, we train the model to approach faces yielding positive interactions and avoid faces yielding negative interactions. Using intracortical electroencephalography (iEEG) data and Representational Dissimilarity Matrices (RDMs), we evaluate the model’s ability to account for neural responses. Our RL model performs at the same level as supervised and unsupervised models, capturing neural responses to complex visual stimuli. The findings suggest that RL models are a promising approach for understanding perception.

**Significance Statement:** Understanding how the brain encodes faces is central to vision science. Existing models rely on supervised learning, which requires ground-truth labels that are often unavailable in real-world settings, or on unsupervised learning, which ignores the role of environmental-feedback in shaping visual representations. We introduce a reinforcement learning (RL) model that learns through environmental feedback, simulating human interactions by associating approaching faces with positive interactions and avoiding faces with negative interactions. Using intracortical electroencephalography (iEEG) data from face-selective regions, we show that an RL model with a variational DenseNet encoder accounts for neural representations comparably to supervised and unsupervised models. Task and architecture jointly shaped representational geometry, highlighting the importance of both learning objective and encoder design. These findings suggest the potential of RL-based approaches to understand neural representations of naturalistic faces.

## 1 Introduction

Faces encode a wealth of socially-relevant information, such as a person’s identity and expressions. What are the computational mechanisms that give rise to neural representations of faces? Recent work modeled neural responses to faces and other objects using supervised deep neural network models that learn by comparing the outputs of the models to ground-truth labels (Khaligh-Razavi and Kriegeskorte, 2014; Cadieu et al., 2014; Schwartz et al., 2023a). These models yielded promising results, but rely on ground-truth labels that are not generally available to human observers. To address this limitation, more recent studies used unsupervised models - that do not need ground-truth labels for training - (Higgins et al., 2021; Zhuang et al., 2021; Konkle and Alvarez, 2022), succeeding at predicting neural responses with accuracy that matches that of supervised models.

Current unsupervised models typically learn face and object representations without taking into consideration the downstream behavioral tasks that observers need to perform. For example, some models are trained to build accurate image reconstructions (Higgins et al., 2021), other models are trained to minimize the dissimilarity between the representation of different images that correspond to a same object (Konkle and Alvarez, 2022). However, several studies indicate that visual representations are also shaped by the tasks animals need to perform, and in particular by the behavioral relevance of distinguishing between different sets of stimuli. For example, when macaques are trained to distinguish between different categories of stimuli to receive a reward, more neurons in inferotemporal cortex become tuned for the stimulus properties that are relevant to make that categorization (even when counterbalancing which properties are relevant across subjects; Sigala and Logothetis, 2002; De Baene et al., 2008). Converging evidence comes from recent work in mice, that used two-photon imaging to show that after category learning ventral visual regions are associated with an increase in the fraction of neurons responsive to the trained task (Goltstein et al., 2021). Together, these studies indicate that the task-relevant feedback an animal receives from the environment contributes to shaping their visual representations.

Unsupervised models of neural responses have difficulty accounting for this kind of dependence of visual representations on the feedback an observer receives from the environment. We sought to address this gap in the case of face perception, using a model trained with a simple approach-avoidance task. While such approach-avoidance does not come close to the full complexity of the behaviors that are informed by face perception in naturalistic settings, representations learned using this task can provide a lower bound of the correspondence that can be obtained between face-selective neural responses and models trained with information that is available in real-world social interactions. To evaluate this, we trained a deep neural network model to approach individuals with whom they have positive interactions, and to avoid individuals with whom they have negative interactions (see Materials and Methods for details of the interactions). We then asked whether this model could account for neural responses to faces recorded by intracranial electrodes.

The model was trained in a simulated environment that included individuals with different identities. In order to account for the variability in the valence of repeated interactions with the same individual, each identity was associated with a probability distribution representing the reward that would be obtained by approaching it. Each identity was also associated with a set of face images varying in viewpoint, illumination, and other properties. At each trial, the model received a face image as input, and chose whether or not to approach the associated identity, receiving a reward as a result of its choice. The model was trained to maximize its reward.

We used representational similarity analysis (Kriegeskorte et al., 2008) to quantify the correspondence between the representations learned by this model and neural responses to faces recorded with intracortical electroencephalography (iEEG). We then compared the correspondence obtained with the model to those obtained with deep supervised and unsupervised models that were built with matching architectures and trained with the same dataset.

## 2 Materials and Methods

### 2.1 Participants

The experimental protocols were approved by the University of Pittsburgh’s Institutional Review Board. Informed consent was acquired from all participants (see also Li et al., 2019; Boring et al., 2021 for details, which used the same data). The sample comprised 11 participants (mean age = 31.8 years, SD = 9.89; 7 females) who underwent intracranial electroencephalography (iEEG) electrode (surface and depth) implantation for seizure onset localization. One participant was removed before performing additional analysis due to noisy data. None of the participants showed evidence of epileptic activity on electrodes located in the ventral and lateral temporal lobes.

### 2.2 Experimental Design

#### 2.2.1 Experiment paradigm

At the beginning of the experiment, all participants completed a functional localizer task that was used to identify face-selective electrodes. Next, they completed the main task. In the main task, each trial consisted of a face image (presented for 1000ms) followed by a 500ms inter-trial interval during which a fixation cross was displayed. Participants were asked to identify the gender of the presented faces, as rapidly and accurately as possible. The face images presented to participants were chosen from the Karolinska Directed Emotional Faces (KDEF) dataset (Lundqvist et al., 1998). This dataset contains 4900 images featuring 70 individuals (50% female) displaying seven distinct expressions from five angles. In this study, we used a subset of expressions, including happy, sad, fearful, angry, and neutral expressions. Each face identity and expression combination was depicted from various angles, including frontal view (0 degrees), left and right 45-degree views, and left and right 90-degree profiles.

In addition, the participants were divided into two subsets, that completed slightly different versions of the experiment (A and B). In version A, participants completed one 200-trial session, viewing a stimuli set comprising eight identities, five expressions, and five viewpoint angles (left/right profile, left/right 45 degree, and frontal). Each stimulus was displayed three times. In version B, participants completed at least two sessions viewing a distinct KDEF subset of 600 images, comprising 40 identities, five expressions, and three viewpoint angles (profile, 45 degree, and frontal). Each stimulus was presented only once.

#### 2.2.2 Data processing

Data was pre-processed at the University of Pittsburgh (Li et al., 2019; Boring et al., 2021). The data encompasses 14 depth and 11 surface electrodes that recorded local field potentials at 1000 Hz. Both types of electrodes had comparable recording site surface areas and recorded similar neural responses.

To extract single-trial potential signals, the raw data underwent band-pass filtering using a fourth order Butterworth filter, with frequencies between 0.2 Hz and 115 Hz preserved. Subsequently, slow and linear drift components, as well as high-frequency noise, were removed. Additionally, a 60 Hz line noise was eliminated using a stop-band encompassing the range of 55-65 Hz. Single-trial potentials (referred to as stP) were time-locked to the onset of each stimulus, with the signal sampled at 1000 Hz. Artifacts were reduced by examining raw data, with no ictal events observed. Trials exceeding 5 standard deviations above the mean maximum amplitude were excluded, as were trials with a difference of *≥*25 *µ*V between consecutive sampling instances, resulting in *<*1% trial removed.

#### 2.2.3 Electrode localization

Electrode location (Fig. 1C) was ascertained using an automated co-registering method (Hermes et al., 2010). Postoperative CT scans were aligned with anatomical MRI scans to sectionalize electrode contacts pre-surgery. Pre- and post-operative imaging scans were also used to localize SEEG electrodes. Face-selective electrodes were determined by analyzing functional localizer data. An electrode was deemed face-selective if it significantly differentiated faces from other object categories (Li et al., 2019; Boring et al., 2021).

**Fig. 1.**
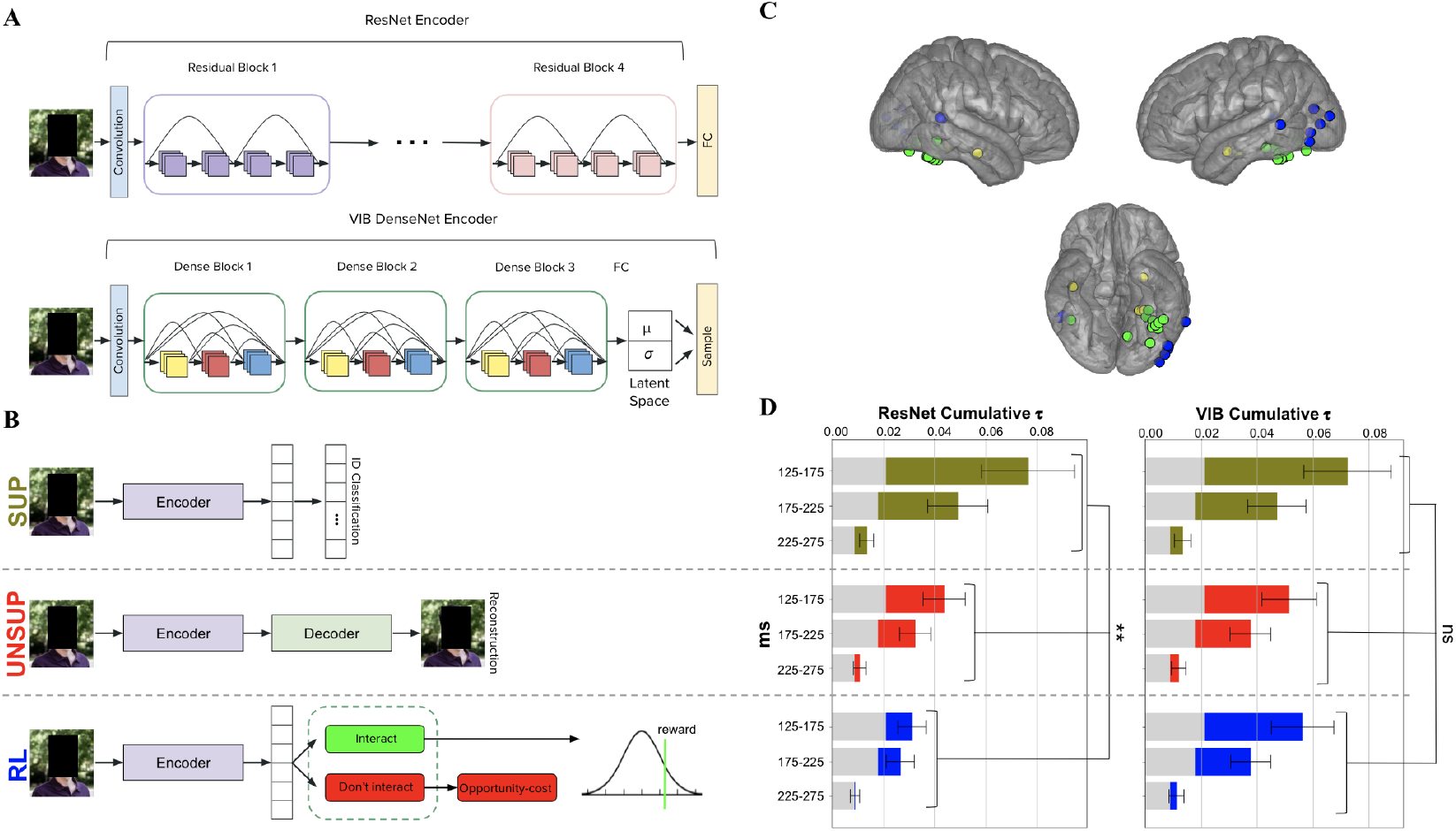
Deep network models and correlations with neural responses. **A**. Two types of encoder architectures. **B**. Neural network architectures for the supervised (SUP), unsupervised (UNSUP), and reinforcement learning (RL) models. **C**. Face-selective electrode locations (*n* = 24). **D**. Cumulative Kendall *τ* correlations between face-selective iEEG RDMs and the RDMs from each model averaged over electrodes (*n* = 24).

##### Face-selective electrode localization

Across the 11 consented participants, a total of 1,079 electrodes were implanted, and 25 electrodes (2.3%) were face selective. Of the 25 electrodes, 12 were in the ventral stream (ventral temporal and occipital cortex anterior to V2), including 10 in fusiform gyrus (localization determined with Neurosynth; Yarkoni et al., 2011). One of these electrodes did not pass the reliability analysis and was excluded from further analyses. The remaining 24 electrodes are shown in Fig. 1C. Eight were in the lateral stream (lateral temporal and lateral occipital cortex anterior to V2, including V3d, V5/MT, and STS); and four were outside these streams (labeled as “other”).

##### Electrode coverage

A total of 24 face-selective electrodes were identified across 10 subjects (after excluding one subject due to noisy data). The number of electrodes contributed by each subject was as follows: P16 (1 electrode), P23 (1 electrode), P27 (5 electrodes), P28 (1 electrode), P30 (1 electrode), P34 (4 electrodes), P36 (3 electrodes), P39 (1 electrode), P41 (6 electrodes), and P47 (1 electrode).

### 2.3 Statistical Analysis

#### 2.3.1 Deep convolutional neural network models

We deployed distinct deep convolutional neural networks (DCNNs), varying in architecture and learning mechanisms, to model the neural data. In order to compare different learning mechanisms while keeping the architectures as comparable as possible, three networks used the same encoder architecture, but varied in their learning mechanism: a supervised model (SUP focused on identity (ID) classification, an unsupervised model (UNSUP) aimed to reconstruct the original image, and a reinforcement learning model (RL) was trained to predict the expected reward associated with interacting with a particular person (identity). We used identity classification rather than gender as the supervised objective because identity is more challenging and more informative for characterizing face-selective representations, whereas gender is typically easier and less informative. These models were constructed based on a residual neural network architecture – ResNet-18 (He et al., 2016; see Fig. 1A), that was also used in recent work on face perception (Schwartz et al., 2023a). In addition to these three models trained on the ResNet encoder architecture, in order to facilitate comparison to different encoder architectures, we also included three models (SUP, UNSUP, and RL) constructed using a combination of densely connected architecture – DenseNet (Huang et al., 2017), and Variational Autoencoders – VAEs (Kingma and Welling, 2013). In addition – since the RL model with this encoder architecture showed competitive correlations with neural responses (see Fig. 1D) – we used this architecture as the basis to test a multi-task model (VIB UNSUP+RL) which combined image reconstruction and reward prediction objectives using this architecture (see Fig. 1A, B & Fig. 5A). The details of the models’ architectures are described in the following section.

#### 2.3.2 DCNN architectures

In a first set of three models, we used a standard residual neural network (ResNet-18) architecture as the encoder (He et al., 2016). In such encoder, input images were processed through a series of convolutional layers with batch normalization and ReLU activation functions (see Table 1).

**Table 1.**
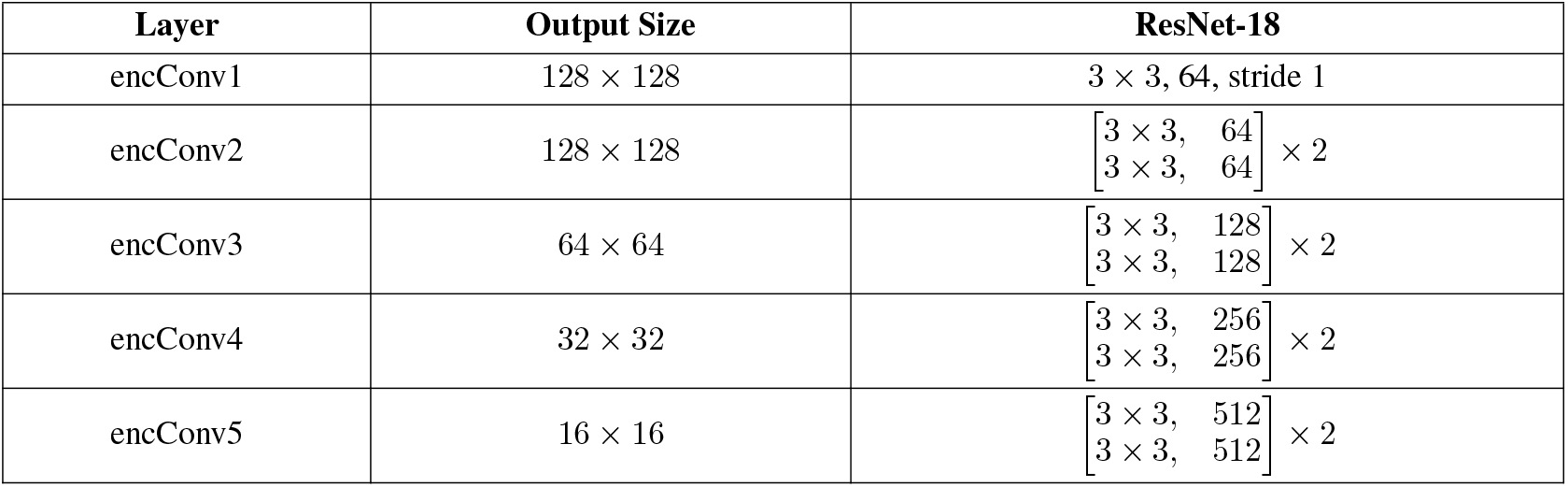
ResNet encoder architecture.

| Layer | Output Size | ResNet-18 |
| --- | --- | --- |
| encConv1 | $128 \times 128$ | $3 \times 3, 64, \text{stride } 1$ |
| encConv2 | $128 \times 128$ | $\begin{bmatrix} 3 \times 3, & 64 \\ 3 \times 3, & 64 \end{bmatrix} \times 2$ |
| encConv3 | $64 \times 64$ | $\begin{bmatrix} 3 \times 3, & 128 \\ 3 \times 3, & 128 \end{bmatrix} \times 2$ |
| encConv4 | $32 \times 32$ | $\begin{bmatrix} 3 \times 3, & 256 \\ 3 \times 3, & 256 \end{bmatrix} \times 2$ |
| encConv5 | $16 \times 16$ | $\begin{bmatrix} 3 \times 3, & 512 \\ 3 \times 3, & 512 \end{bmatrix} \times 2$ |

While we used the same encoder architecture for the three models, each model was based on a different decoder architecture that varied depending on the task the model needed to perform. Each decoder operated on the representations derived from the encoder module. For the ResNet SUP model, a classifier was utilized to perform identity classification based on the feature representations. The classifier consisted of one fully connected layer (see Table 2). For the ResNet UNSUP model, the decoder component first generated a vector of means (*µ*) and a vector of standard deviations (*σ*) using fully connected layers. These two parameters are essential for the image reconstruction process (see details in the following paragraph on VAEs). The rest of the decoder component mirrored a symmetric structure to that of the encoder. Transpose convolutional layers were utilized for upsampling, ensuring the preservation of spatial information (see Table 3). Finally, the decoder component of the ResNet RL model was based on a network that generated as output the predicted reward that would be derived from interacting with a face. Based on this output, the network then chose whether or not to interact with the face, and a loss was computed based on the chosen interaction (or lack thereof). The details of the loss function and training procedure are described in the following section. The network utilized five fully connected layers, with the first four including ReLU activation functions (see Table 4).

**Table 2.** ResNet SUP model: decoder architecture.

| Layer | In features | Out features | Input Source |
| --- | --- | --- | --- |
| layer1 | 512 | 1503 | encConv5 |

**Table 3.**
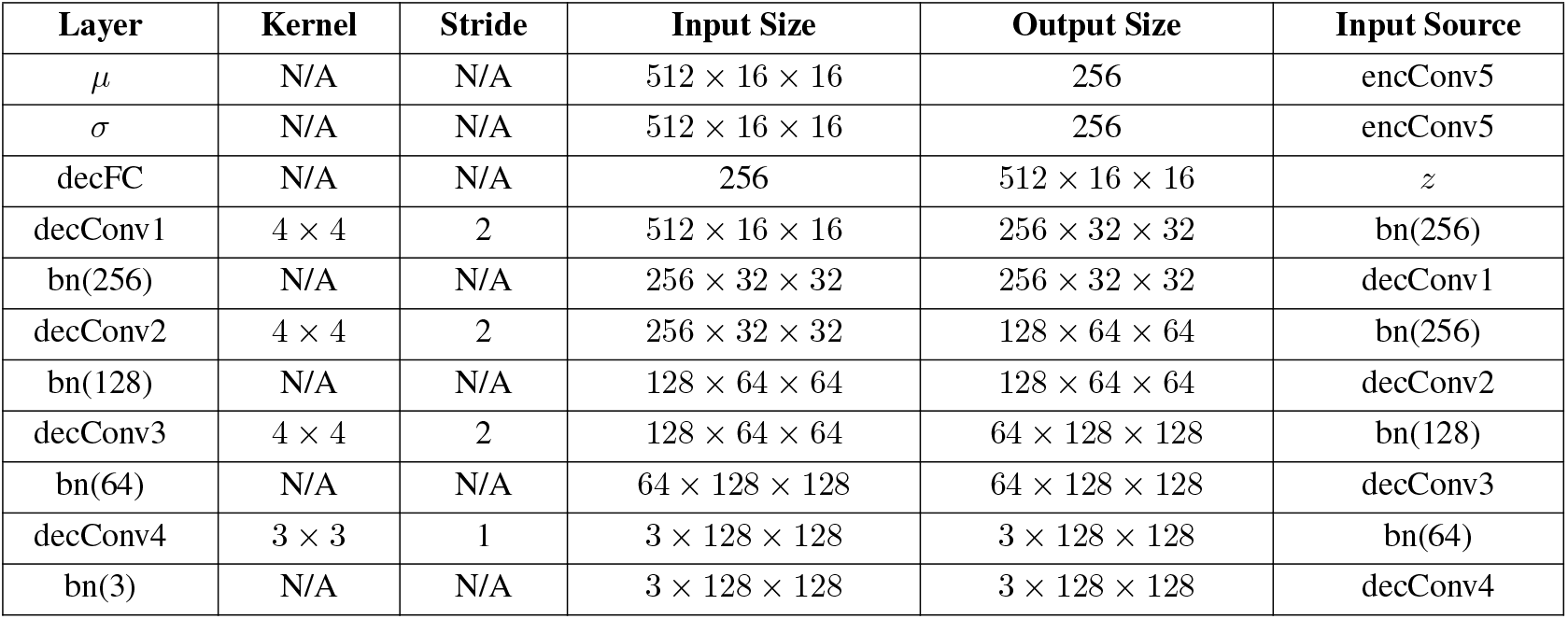
ResNet UNSUP model: decoder architecture. bn: batch normalization.

**Table 4.** ResNet RL model: decoder architecture.

| Layer | In features | Out features | Input Source |
| --- | --- | --- | --- |
| layer1 | 512 | 1024 | FC |
| relu1 | 1024 | 1024 | layer1 |
| layer2 | 1024 | 512 | relu1 |
| relu2 | 512 | 512 | layer2 |
| layer3 | 512 | 256 | relu2 |
| relu3 | 256 | 256 | layer3 |
| layer4 | 256 | 128 | relu3 |
| relu4 | 128 | 128 | layer4 |
| layer5 | 128 | 1 | relu4 |

In a second set of three additional models, we used an encoder architecture that combines the inherent strengths of DenseNet (Huang et al., 2017) and variational autoencoders – VAEs (Kingma and Welling, 2013). The inclusion of DenseNet-like skip-connections in the encoder improved identity classification performance based on the latent representations generated by the encoder, making it possible to compare the learning mechanisms building on the same encoder architecture.

In the encoder, input images were processed through a series of convolutional layers with batch normalization and ReLU activation functions (see Table 5). The encoder consists of three dense blocks, each comprising three convolutional layers. Within each dense block, feature maps from different layers are concatenated along the channels dimension. This design choice promotes feature reuse and alleviates the vanishing gradient problem. After the dense blocks, the resulting feature maps are then fed into fully connected layers, which generate a vector of means (*µ*) and a vector of standard deviations (*σ*) of the latent space (as in a typical variational autoencoder; Kingma and Welling, 2013). These vectors represent the parameters of a Gaussian distribution associated with a particular input stimulus. To incorporate stochasticity in the model and enable probabilistic sampling, the reparameterization trick is applied. This trick involves sampling latent variables by adding random noise *ϵ* drawn from a Gaussian distribution with zero mean and unit standard deviation. The latent variable *z* is computed as *z* = *µ* + *σ · ϵ*. Then, *z* is used as the input to the decoder network, which varies depending on the task (details are provided in the following paragraph). This stochastic layer introduces regularization and enables the use of the network as a generative model (Kingma and Welling, 2013).

**Table 5.** VIB DenseNet encoder architecture. bn: batch normalization.

| Block | Layer | Kernel | Stride | Input Size | Output Size | Input Source |
| --- | --- | --- | --- | --- | --- | --- |
| Conv 1 | bn1(3) | N/A | N/A | $3 \times 128 \times 128$ | $3 \times 128 \times 128$ | Images |
| | encConv1 | $4 \times 4$ | 2 | $3 \times 128 \times 128$ | $64 \times 64 \times 64$ | bn1 |
| | bn2(64) | N/A | N/A | $64 \times 64 \times 64$ | $64 \times 64 \times 64$ | encConv1 |
| Dense 1 | encConv2 | $3 \times 3$ | 1 | $64 \times 64 \times 64$ | $64 \times 64 \times 64$ | bn2 |
| | bn3(128) | N/A | N/A | $128 \times 64 \times 64$ | $128 \times 64 \times 64$ | encConv1, encConv2 |
| | encConv3 | $3 \times 3$ | 1 | $128 \times 64 \times 64$ | $64 \times 64 \times 64$ | bn3 |
| | bn4(192) | N/A | N/A | $192 \times 64 \times 64$ | $192 \times 64 \times 64$ | encConv1, encConv2, encConv3 |
| | encConv4 | $3 \times 3$ | 1 | $192 \times 64 \times 64$ | $64 \times 64 \times 64$ | bn4 |
| | bn5(64) | N/A | N/A | $64 \times 64 \times 64$ | $64 \times 64 \times 64$ | encConv4 |
| | AvgPool2d(2) | N/A | N/A | $64 \times 64 \times 64$ | $64 \times 32 \times 32$ | bn5 |
| Dense 2 | encConv5 | $3 \times 3$ | 1 | $64 \times 32 \times 32$ | $64 \times 32 \times 32$ | AvgPool2d(2) |
| | bn6(128) | N/A | N/A | $128 \times 32 \times 32$ | $128 \times 32 \times 32$ | encConv4, encConv5 |
| | encConv6 | $3 \times 3$ | 1 | $128 \times 32 \times 32$ | $64 \times 32 \times 32$ | bn6 |
| | bn7(192) | N/A | N/A | $192 \times 32 \times 32$ | $192 \times 32 \times 32$ | encConv4, encConv5, encConv6 |
| | encConv7 | $3 \times 3$ | 1 | $192 \times 32 \times 32$ | $64 \times 32 \times 32$ | bn7 |
| | bn8(64) | N/A | N/A | $64 \times 32 \times 32$ | $64 \times 32 \times 32$ | encConv7 |
| Dense 3 | encConv8 | $3 \times 3$ | 1 | $64 \times 32 \times 32$ | $64 \times 32 \times 32$ | bn8 |
| | bn9(128) | N/A | N/A | $128 \times 32 \times 32$ | $128 \times 32 \times 32$ | encConv7, encConv8 |
| | encConv9 | $3 \times 3$ | 1 | $128 \times 32 \times 32$ | $64 \times 32 \times 32$ | bn9 |
| | bn10(192) | N/A | N/A | $192 \times 32 \times 32$ | $192 \times 32 \times 32$ | encConv7, encConv8, encConv9 |
| | encConv10 | $3 \times 3$ | 1 | $192 \times 32 \times 32$ | $64 \times 32 \times 32$ | bn10 |
| | bn11(64) | N/A | N/A | $64 \times 32 \times 32$ | $64 \times 32 \times 32$ | encConv10 |
| FC | $\mu$ | N/A | N/A | $64 \times 32 \times 32$ | 2048 | bn11 |
| | $\sigma$ | N/A | N/A | $64 \times 32 \times 32$ | 2048 | bn11 |

As in the case of the models using the ResNet encoder, for the VIB DenseNet models, each model was based on a different decoder architecture that varied depending on the task. Each decoder operated on the latent representations derived from the encoder module. For the VIB SUP model, fully connected layers were employed to generate the mean (*µ*) and standard deviation (*σ*) for the latent space after the encoder. Then, a classifier was utilized to perform identity classification based on the feature representations. The classifier consisted of two fully connected layers, with the first layer including a ReLU activation function (see Table 6). For the VIB UNSUP model, the decoder component mirrored a symmetric structure to that of the encoder. Transpose convolutional layers were utilized for upsampling, ensuring the preservation of spatial information (see Table 7). Finally, the decoder component of the VIB RL model was based on a network built to predict reward using the latent variable *z* from the encoder as input. It utilized five fully connected layers, with the first four including ReLU activation functions (see Table 8).

**Table 6.** VIB SUP model: decoder architecture.

| Layer | In features | Out features | Input Source |
| --- | --- | --- | --- |
| layer1 | 2048 | 2048 | FC |
| relu1 | 2048 | 2048 | layer1 |
| layer2 | 2048 | 1503 | relu1 |

**Table 7.**
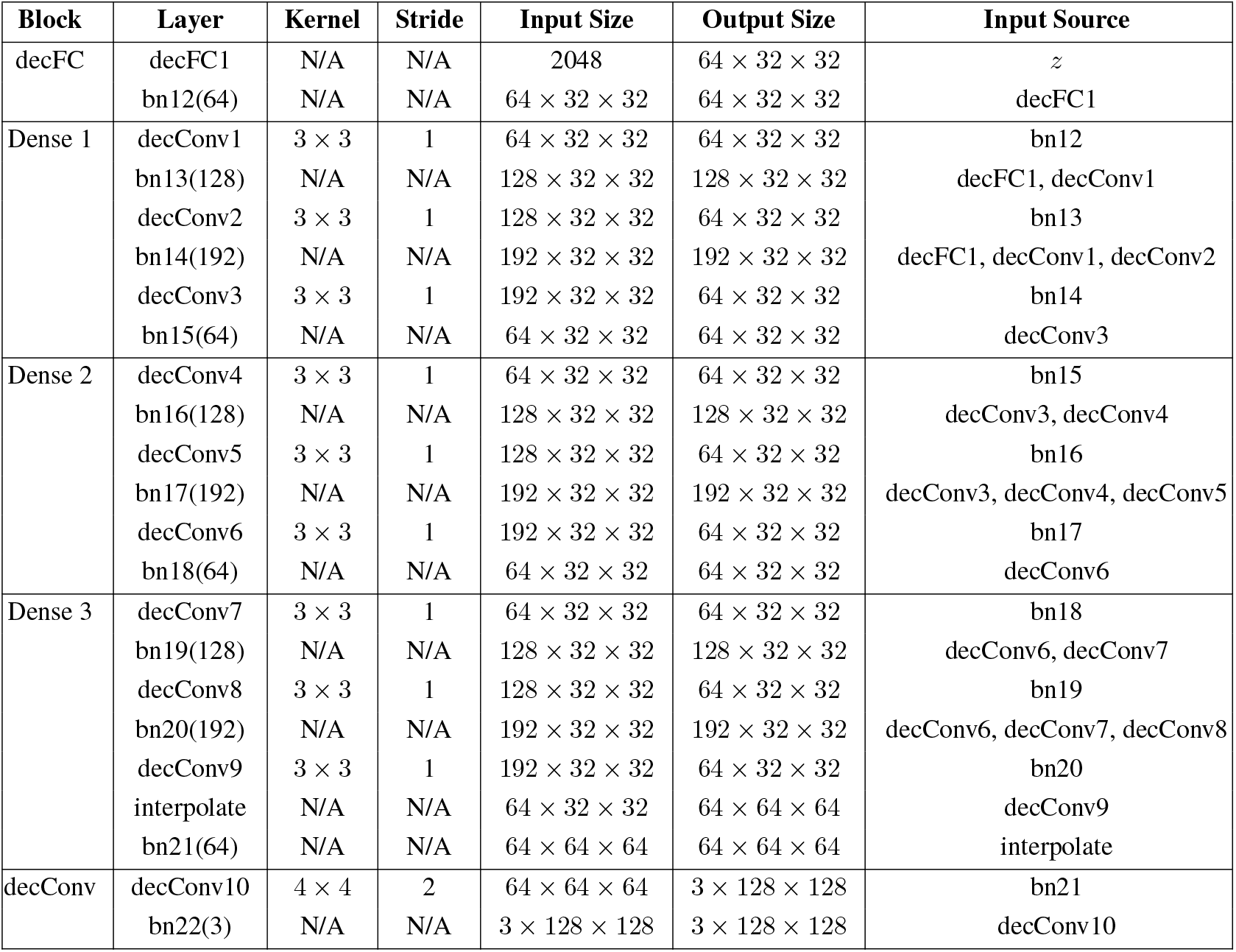
VIB UNSUP model: decoder architecture. bn: batch normalization.

**Table 8.** VIB RL model: decoder architecture.

| Layer | In features | Out features | Input Source |
| --- | --- | --- | --- |
| layer1 | 2048 | 1024 | FC |
| relu1 | 1024 | 1024 | layer1 |
| layer2 | 1024 | 512 | relu1 |
| relu2 | 512 | 512 | layer2 |
| layer3 | 512 | 256 | relu2 |
| relu3 | 256 | 256 | layer3 |
| layer4 | 256 | 128 | relu3 |
| relu4 | 128 | 128 | layer4 |
| layer5 | 128 | 1 | relu4 |

#### 2.3.3 Loss functions

Each of the models had a corresponding loss function designed for the task the model needed to complete. The SUP models used a standard Cross-Entropy loss for classification. The UNSUP models used a reconstruction loss, and the RL models used an RL loss. In addition, all of the models using the VIB encoder architecture have a KL-divergence loss component added to the overall loss function. The VIB UNSUP+RL model used the following loss function:

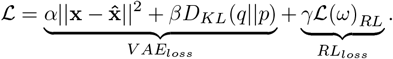

Where x is an image of a person’s face, *α* = 4000, *β* = 0.0000001, and *γ* = 1.5.

The loss ℒ (*ω*)_*RL*_is a reinforcement learning loss constructed as follows. Each person identity *i* is associated with a reward distribution 𝒩 (*µ*_*i*_, *σ*_*i*_). We modeled the reward distribution per identity as Gaussian to capture stochastic feedback expected from natural social interactions. The mean *µ*_*i*_ encodes identity-specific expected value, while *σ*_*i*_ captures uncertainty. The network *h* produces as output the expected reward 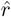. The model chooses to interact with the person in the image with probability 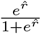. If the model chooses to interact with the person (*ω* = 1), it obtains a reward *r* ∼ 𝒩(*µ*_*i*_, *σ*_*i*_), and computes a ‘prediction error’ loss given by the square of the difference between the obtained reward and the expected reward. Instead, if the model chooses to not interact with the person (*ω* = 0), it computes an ‘opportunity cost’ loss associated with not interacting.

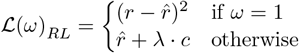

The ‘opportunity cost’ is computed with:

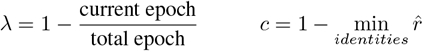

Therefore, at the start of training, the opportunity cost was positive and greater or equal to 1 for all identities, promoting exploration. This choice was inspired by the idea of the optimistic initial value in reinforcement learning. As training progressed, and the model’s predictions of the rewards that would result from interacting with different face images improved, the decay of *λ* led the opportunity cost to converge to the predicted rewards 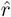Note that the opportunity cost loss is essential to prevent the network’s outputs to collapse to small numbers. In fact, in its absence, the network learns to predict small values of 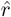 this prevents the model from incurring in the prediction error loss, because the prediction error loss is incurred by the model only if it chooses to interact, and the probability of interacting is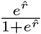which is low for low 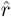values.

#### 2.3.4 Training and testing

All models were trained using the CelebA dataset (Liu et al., 2018) consisting of over 300,000 images. In order to match the size of the dataset with the recent work (Schwartz et al., 2023a) to facilitate comparison between studies, a subset of CelebA was utilized. This subset contained 28,709 training images and an additional 3,589 labeled testing images, featuring a total of 1,503 unique identities. Random selection was employed to ensure a minimum of 20 images per identity. All images were uniformly resized to dimensions of 128 × 128 pixels and RGB scale.

Following the training, the ResNet SUP and VIB SUP networks were tested on their ability to perform identity recognition using the CelebA dataset. The testing extracts the output from each network for every image in the dataset. ResNet SUP model was able to recognize identity, achieving an accuracy of 22.76 % on a left-out subset of CelebA (chance level = 0.07%). VIB SUP model was able to recognize identity, achieving an accuracy of 18.5% on a left-out subset of CelebA (the classification accuracy was 6.88% without the DenseNet architecture). Other results about model performances will be described in the “Results” section below.

#### 2.3.5 Temporal localizer

In order to compare the representations in DCNNs to neural representations, we first aimed to identify temporal windows during which face-selective electrodes yielded the most consistent responses. The data time-series was segmented into successive, non-overlapping 50*ms* time windows, starting from 25*ms* pre-stimulus onset up to 525*ms* post-onset. We correlated all instances of a stimulus’s neural response within each time window and calculated an average correlation across all time windows for that stimulus. The overall average correlation was then subtracted from the time-window-specific correlations. We then conducted a paired t-test to distinguish time windows with above-average test-retest reliability (p *<* 0.05). Following Bonferroni correction for multiple comparisons, one electrode was omitted from the RDM analysis due to a lack of reliable response time windows (p *>* 0.05).

#### 2.3.6 Computing representational similarity analysis for neural data

Aiming to preserve as much data as possible, we initially analyzed all face-selective electrodes, including those exposed to every stimulus once. We calculated Representational Dissimilarity Matrices (RDMs) for three temporal windows (125*ms* - 175*ms*, 175*ms* - 225*ms*, 225*ms* - 275*ms*) based on previous research on visual face perception’s timing. A 50-dimensional vector was derived for each temporal window per electrode, with each dimension representing the response level per millisecond of the 50*ms* window. Using correlation distance, we measured the dissimilarity between each pair of stimulus response patterns to obtain neural RDMs.

Following this procedure, RDMs of dimensions 200×200 were generated for Experiment A (5 expressions times 5 viewpoints times 8 identities), while RDMs of dimensions 600×600 were obtained for Experiment B (5 expressions times 3 viewpoints times 40 identities). It is important to note that the sizes of the RDMs differed between the two experiments because different subsets of KDEF images were used in Experiment A and Experiment B, as described in the “Experimental paradigm” section. Specifically, in Experiment B, the information regarding viewpoint only included the viewpoint angle, without distinguishing between left and right viewpoints. Consequently, the feature vectors for the left and right viewpoints were averaged, resulting in the averaging of left and right profile views as well as left and right 45 degree views.

#### 2.3.7 Computing representational similarity for DCNNs

To study the similarity between the representations in different networks, we used representational similarity analysis (RSA). Specifically, for the VIB DenseNet encoder architecture, features from the Variational Information Bottleneck (VIB) representations were extracted; for the ResNet encoder architecture, features from the final layer in the last residual blocks were extracted. For each of the models, we calculated representational dissimilarity matrices (RDMs) using a three-step procedure that has been used in previous studies (Schwartz et al., 2023a). First, feature vectors were extracted for all KDEF images employed in the experiment. Subsequently, the feature vectors were mean-centered by subtracting the mean feature vector across all KDEF images. Finally, for each pair of images, the correlation distance between their mean-centered feature vectors was computed using Pearson’s correlation coefficient (*r*), and the correlation distance was defined as 1 *™ r*.

#### 2.3.8 Comparing RDMs

Once the RDMs for each of the electrodes and time windows were computed, as well as the RDMs for the different computational models, the similarity between neural RDMs and model RDMs was calculated using the Kendall *τ* rank correlation coefficient (*τ*_*B*_) (Fig. 1D). In addition, Kendall’s *τ*_*B*_ was also used to calculate the similarity between the RDMs of different computational models (Fig. 2B).

**Fig. 2.**
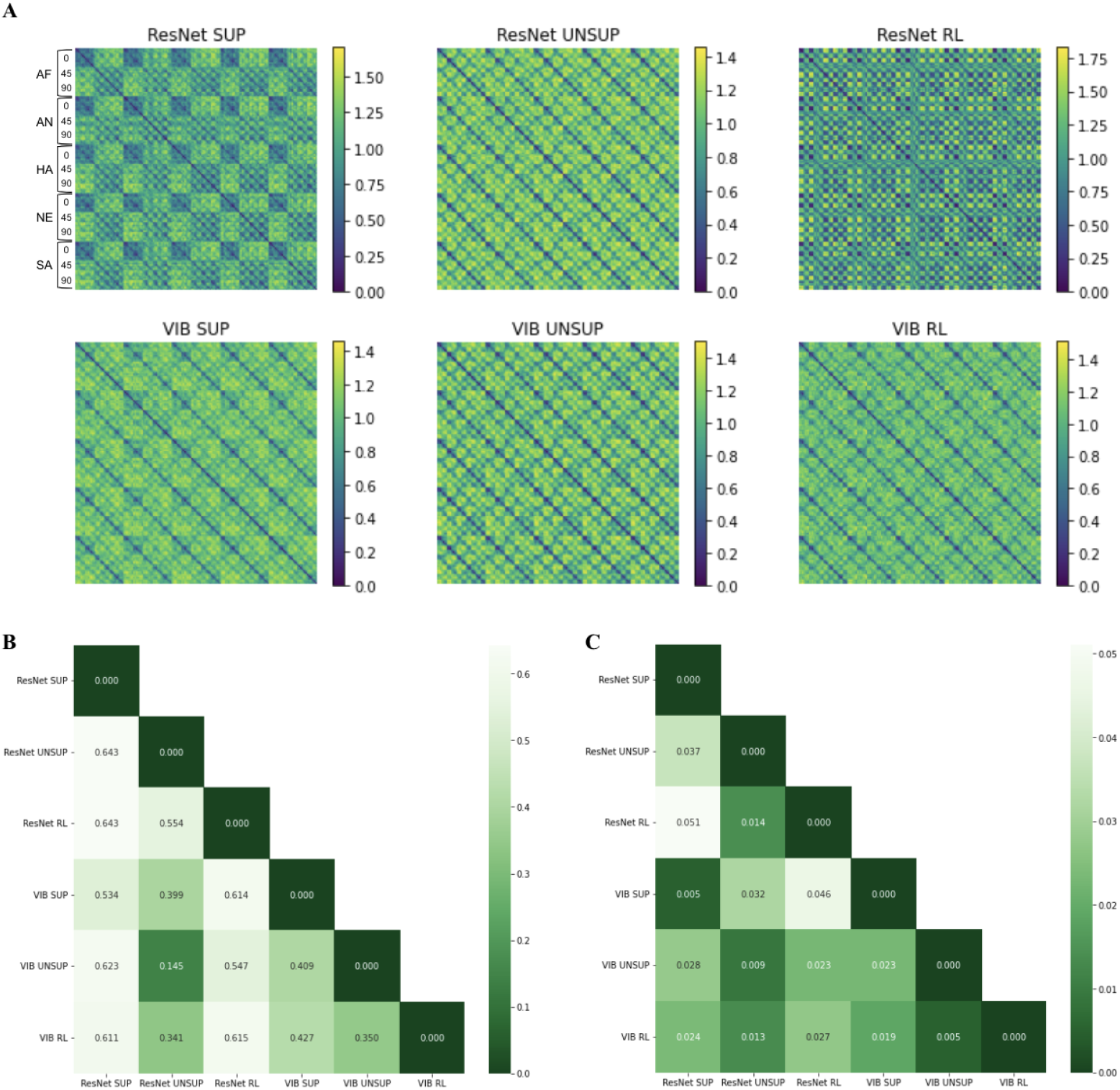
Model comparisons. **A**. RDMs of the latent representations of all models (based on KDEF images used in version A of the experiment). **B**. Kendall’s *τ* ranking distance between the RDMs of the different models. **C**. Euclidean distance between the models’ match with neural responses. The match was computed as cumulative Kendall *τ* values between model RDMs and neural RDMs.

## 3 Results

### 3.1 Deep neural network models of face perception

Six deep neural network models of face perception were trained using a subset of the CelebA dataset (Liu et al., 2018). In order to evaluate models that learn based on feedback from the environment, we implemented a deep convolutional neural network (DCNN) trained with the approach-avoidance task described in the Introduction. Because the model was trained to maximize its reward through interactions with its environment, it is a type of reinforcement learning model, and we will refer to it as such throughout the article (“RL model”). However, it should not be confused with other kinds of reinforcement learning models, such as deep Q-learning, that use different training mechanisms.

In addition to the RL model, we implemented two comparison DCNNs: one trained with a supervised task (“SUP model”), and one trained with an unsupervised task (“UNSUP model”). Given an image of a person’s face as input, the SUP model was trained to classify the identity of that face. The UNSUP model was trained to reconstruct the input image as accurately as possible. The RL model was trained to predict the reward that would result from interacting with the identity associated with that face image, and to approach identities yielding positive rewards. The RL model differs from the SUP model in two important ways. First, in the RL model, during training, different identities can be associated with similar rewards. By contrast, in the SUP models, different identities are always associated with different labels. Second, for the RL model there is a trade-off between acquiring information and receiving reward: choosing to approach identities associated with negative rewards provides information, but it also has a cost. Supervised and unsupervised DCNNs have been used to model neural responses to faces in previous work (Tsantani et al., 2021; Schwartz et al., 2023a; Higgins et al., 2021). Here, we aimed to test whether RL models, that do not have access to the ground truth labels but can receive feedback from the environment, can also provide comparable results when they are equipped with the same encoder architecture.

The performance of a model and its correspondence with neural responses can depend not only on the task the model is trained with (e.g. SUP, UNSUP, or RL), but also on the model’s architecture. In order to evaluate the impact of the architecture, we tested models built on two different encoder architectures (Fig. 1A). ResNet-18 was selected as one of the encoder architectures because ResNet architectures are widely used in neuroscience (Wen et al., 2018; Schwartz et al., 2023a; Dobs et al., 2023); therefore, facilitating comparison with previous studies. A modified version of a variational model was selected as the other: this choice was motivated by recent results showing that variational architectures learn factors that show correspondence with the tuning properties of neurons in inferotemporal cortex (Higgins et al., 2021). The original version of this variational architecture did not perform well at the supervised task; therefore, we modified it with the addition of dense connections (Huang et al., 2017); this change improved its performance at supervised identity classification. For each encoder, we used three different decoders: each decoder was tailored to the type of output the model needed to produce (Fig. 1B). While some of the models shared the same encoder architecture, each model was individually trained end-to-end to perform one task only. Thus, even models with the same encoder architecture ultimately learned different encoder weights.

First, we ensured that the RL models successfully learned to interact with the faces yielding positive rewards. Indeed, there were significant increase in rewards after training the RL models. We trained the RL models ten times using both ResNet and VIB DenseNet encoder. For the ResNet RL model, the average reward before training was −0.255 (SEM = 0.033), and the average reward after training was 12.604 (SEM = 0.047). For the VIB RL model, the average reward before training was 0.500 (SEM = 0.041), and the average reward after training was 10.671 (SEM = 0.209). The SUP and UNSUP models also learned successfully to perform their tasks – details of their loss function can be found in the Supplemental Material.

### 3.2 Comparing face-selective neural responses to deep networks trained with feedback from the environment

To quantify the correspondence between models and neural responses, we collected intracortical electroencephalography (iEEG) data from 11 participants while they observed images of faces varying in identity, expression, and viewpoint (from the Karolinska Directed Emotional Faces dataset: KDEF; Lundqvist et al., 1998). Participants were asked to classify the gender of the faces (see Rossion et al., 2011; Ghuman et al., 2014; Li et al., 2019; Boring et al., 2021). We analyzed 24 face-selective electrodes from 10 participants (for details, see Materials and Methods and Fig. 1C).

Having identified the face-selective electrodes, we next sought to compare representations measured by these electrodes to representations in the DCNN models. Because neural responses change dynamically over time, and their timecourse can vary from region to region, we compared models to neural responses separately for different time windows (see Materials and Methods and Schwartz et al., 2023a for more details), consistent with previous research on the timing of face processing in the human brain (Rossion et al., 2011). For each electrode and time window, we computed neural representational dissimilarity matrices (RDMs). We also computed RDMs obtained from the DCNN models, by feeding as input to the DCNNs the same KDEF images that were shown to the participants. Finally, Kendall’s *τ* was used to compare the neural RDMs to the RDMs obtained from the DCNNs (Fig. 1D). For all six models we trained, the DCNN RDMs were computed based on features extracted from the final layer of the models’ encoders.

We first tested the correspondence between the RL models’ RDMs, and the RDMs obtained from neural responses in face selective electrodes. Kendall *τ* correlations between the model RDMs and the neural RDMs were positive (Fig. 1D). The Kendall *τ* correlations between model RDMs and neural RDMs were different for the SUP, UNSUP, and RL models using the ResNet encoder. The model trained with the RL task showed lower correlations with neural responses. The main effect of model type was significant (*F* (2, 213) = 5.626, *p* = 0.004, see Fig. 1D). In contrast, when using the VIB DenseNet encoder, the Kendall *τ* correlations were similar across all models, with a non-significant main effect for the model type (*F* (2, 213) = 1.098, *p* = 0.336, see Fig. 1D). This indicates that, when using the VIB DenseNet encoder, the RL model achieved comparable performance to the SUP and UNSUP models. Additionally, to further investigate the influence of architecture within the RL training framework, we directly compared the correlations of the VIB RL and ResNet RL models with neural RDMs using a paired *t*-test across participants. For this analysis, the correlation values for each participant were averaged across all face-selective electrodes and time windows. The VIB RL model showed significantly higher correlations with neural RDMs compared to the ResNet RL model (*t*(9) = 2.565, *p* = 0.030). Since this analysis was a pre-specified comparison limited to the RL models, we did not apply a correction for multiple comparisons.

The correspondence between the model RDMs and neural RDMs was highest in the first time window (125*ms* - 175*ms*), and decreased in subsequent time windows (Fig. 1D). This decrease was consistent across all six models (ResNet SUP, ResNet UNSUP, ResNet RL, VIB SUP, VIB UNSUP, and VIB RL). In order to determine whether the models captured the similarity between neural responses to different images more accurately than simple pixel-based metrics, we additionally performed a comparison between the neural RDMs and RDMs computed based on the Pearson’s correlation distance between the pixel values for each pair of images. The results (represented by the gray bars in Fig. 1D) showed considerably lower Kendall *τ* values when using the pixel-based similarity compared with the six deep network models.

In sum, three main findings emerged from the analyses: first, features from deep network models captured neural representations more accurately than pixel-based metrics, second, the correspondence between deep network models and neural responses was highest in the 125*ms* - 175*ms* time window, third, most importantly, reinforcement learning models with a VIB DenseNet encoder architecture performed comparably to supervised and unsupervised models at accounting for neural responses in face-selective electrodes.

### 3.3 Studying the impact of architecture and task mechanisms on representational geometry

Since we trained models with the same encoder architecture using different tasks, as well as models that differ in terms of their encoder architecture but share the same task, we were able to study the impact of the architecture and of the task on the models’ representational geometry, and on their correspondence with neural responses. To do this, we first computed representational dissimilarity matrices for each model (Fig. 2A), and then the dissimilarity between the dissimilarity matrices themselves (Fig. 2B). In addition, we computed the dissimilarity between models in terms of how they correspond to neural responses. Specifically, for each model, we computed a vector of Kendall’s *τ* correlations between the model RDM and neural RDMs across all face-selective electrodes and time windows. These vectors reflect the extent to which each model matches the representational structure of neural data. We then computed the euclidean distance between these vectors (Fig. 2C) to quantify how similarly two models explain the neural data. The choice of euclidean distance is motivated by the observation that the magnitude of the Kendall *τ* values is important to establish the dissimilarity between models (in addition to the pattern of *τ* values across electrodes), and dissimilarity metrics based on correlation or cosine distance would not take into account such differences in magnitude. Note that two models that have intermediate levels of dissimilarity to each other (i.e. models that correlate with Kendall *τ* values smaller than 1) might be similar in terms of their correspondence to neural responses if the correspondence is driven by overlapping parts of the models’ variance, alternatively, they might be different in their correspondence to neural responses if it is driven by non-overlapping parts of their variance. Thus, the direct similarity between two models might be different from the similarity in the extent to which those models match with neural responses.

The RDMs obtained from the ResNet SUP and ResNet RL models were different from each other and from those obtained with all other models. Conversely, the ResNet UNSUP model, despite having the same encoder architecture, learned more similar representations to the VIB models than to the other ResNet models (Fig. 2B). These patterns indicate that the task used to train the models had an impact on the learned representations within a given encoder architecture, but models trained with same tasks need not be most similar across architectures. In addition, model architecture had an impact on the results as well. Models using the VIB DenseNet encoder showed more similar representations to each other, despite the differences in the decoder architectures and the tasks (Fig. 2B).

In terms of the models’ ability to capture neural responses, task was an important driver of the similarity between models. The SUP models were similar to each other, despite their different encoder architectures (ResNet vs VIB DenseNet). This was also the case for the two UNSUP models, but not for the RL models: there was a greater difference between the ResNet RL model and the VIB RL model in terms of their ability to account for neural RDMs. In addition, the ResNet RL model was very different from the SUP models in how it accounted for neural responses, regardless of the encoder architecture being used (Fig. 2C).

### 3.4 Using deep networks to study the differences between ventral and lateral face-selective regions

After comparing face-selective neural responses to deep networks trained with different learning mechanisms, we studied whether these models provide insights into the distinct functional roles of face-selective representations encoded in ventral temporal regions and lateral temporal regions. The ventral and lateral streams have been hypothesized to be specialized respectively for the recognition of face identity and facial expressions. However, this view has been recently challenged (Duchaine and Yovel, 2015; Li et al., 2019; Schwartz et al., 2023b), raising the question of what are the functional roles of the two streams. Instead of differing based on their role for identity and expressions, representations in ventral and lateral regions might differ in the extent to which they are shaped by feedback from the environment. Feedback-dependent neural tuning has been reported in inferior temporal cortex (Sigala and Logothetis, 2002), and thus ventral regions might be better captured by the model trained with the approach-avoidance task. Alternatively, all models could account similarly well for responses in ventral and lateral regions, and these regions might differ along other dimensions. To determine this, we tested whether the multivariate patterns of correspondences between neural responses and multiple different model types provide information that differentiates between ventral and lateral electrodes.

Together, a total of six models were evaluated. Since for each model we computed the correspondence with neural responses in three time windows, each electrode was associated with 18 Kendall *τ* values (one for each of the 6 models and 3 time windows). We used a logistic regression to classify between ventral and lateral electrodes based on these 18-dimensional patterns of Kendall *τ* values, using a leave-one-electrode-out cross-validation procedure. This resulted in a classification accuracy of 75% – the datapoints and the separating hyperplane are shown in Fig. 3A, projected on the first three principal components for visualization purposes. This result shows that it was possible to distinguish between ventral and lateral electrodes based on the 18-dimensional patterns of correspondences across models and time windows.

**Fig. 3.**
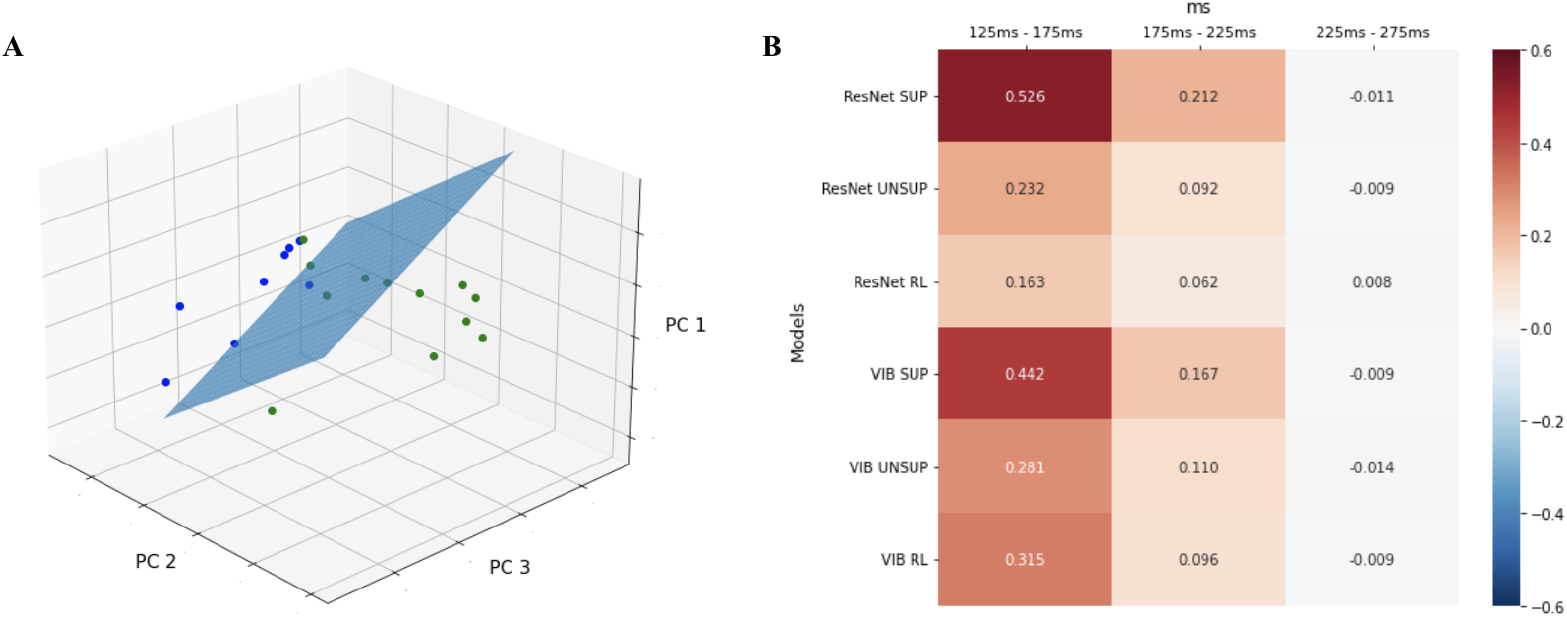
Differences between ventral and lateral electrodes. **A**. Separating hyperplane for the classification of ventral (green) vs lateral (blue) electrodes based on Kendall *τ* values, plotted in the space spanned by the first three principal components. **B**. Coefficients (*β*) of the logistic regression.

To investigate more precisely what information contributes to distinguishing between ventral and lateral electrodes, we extracted the *β* values from the logistic regression (Fig. 3B). A positive *β* value for a particular model and time window means that greater Kendall *τ* values between that model and neural responses in that time window in a particular electrode indicate that the electrode is in ventral temporal cortex. Conversely, if the *β* value for a model and time window is negative, greater correspondence between the model and neural responses in that window indicates that the electrode is in lateral temporal cortex. Therefore, if *β* values exhibit significant variation across model types, it would show that different models contribute differently to distinguishing between ventral and lateral responses. Following the same logic, significant variation in *β* values across time windows would indicate that different time windows contribute differently to distinguishing between ventral and lateral responses.

In order to quantify these effects, we performed a three-way ANOVA testing the variation in *β* values across model types, time windows, and brain regions. The three-way interaction was non-significant (*F* (10, 324) = 0.744, *p* = 0.682), as were the interaction between model type and time window (*F* (10, 324) = 1.137, *p* = 0.333) and the interaction between model type and brain region (*F* (5, 324) = 1.746, *p* = 0.124). The interaction between time window and brain region (ventral vs lateral) was significant (*F* (2, 324) = 24.897, *p <* 0.001). This indicates that across time, ventral and lateral electrodes varied in terms of their correspondence to the models. All main effects were significant (model type: *F* (5, 324) = 4.874, *p <* 0.001; time window: *F* (2, 324) = 48.696, *p <* 0.001; brain region: *F* (1, 324) = 55.974, *p <* 0.001).

In the first two time windows all models had positive *β* values, indicating a greater correspondence with ventral RDMs. What drives this effect? Previous work has demonstrated that lateral regions exhibit stronger responses to dynamic stimuli (Pitcher et al., 2019). We hypothesize that experiments using dynamic stimuli might lead to more reliable responses in lateral regions, and that relatively lower Kendall *τ* values with lateral electrodes might be due to the lack of dynamic information in our stimuli. In the third time window *β* values dropped to near-zero. This drop is likely due to a floor effect: Kendall *τ* values between the models and neural responses were overall low in the third time window.

### 3.5 Studying the unique contributions of different models

Different models can account for distinct or overlapping variance in the neural RDMs. To investigate the degree of overlap between different models, and their unique contributions to capturing the representational structure of neural data, we employed a semi-partial Kendall *τ* analysis. This approach allowed us to isolate and quantify the correlation between each model and neural RDMs, controlling for each other model one at a time. The semipartial Kendall *τ* analysis thus offers a more nuanced understanding of how each model aligns with, or diverges from, the neural representations observed in the iEEG data.

To compute semi-partial Kendall *τ*, we regressed out the RDMs of the computational models in the columns of Fig. 4 from those in the rows, and correlated the residuals with the neural data. The results showed that the supervised models (ResNet SUP and VIB SUP) contain a considerable amount of unique information compared to models trained with the other tasks. This finding underscored the ability of SUP models to explain neural variance that remain unaccounted for by other models. Furthermore, the unique contribution of the models using VIB DenseNet encoder architecture also suggested their ability to capture unique neural activity. Together, these findings pointed to the ability of supervised models and VIB DenseNet architectures to explain neural responses that other models and architectures could not.

**Fig. 4.**
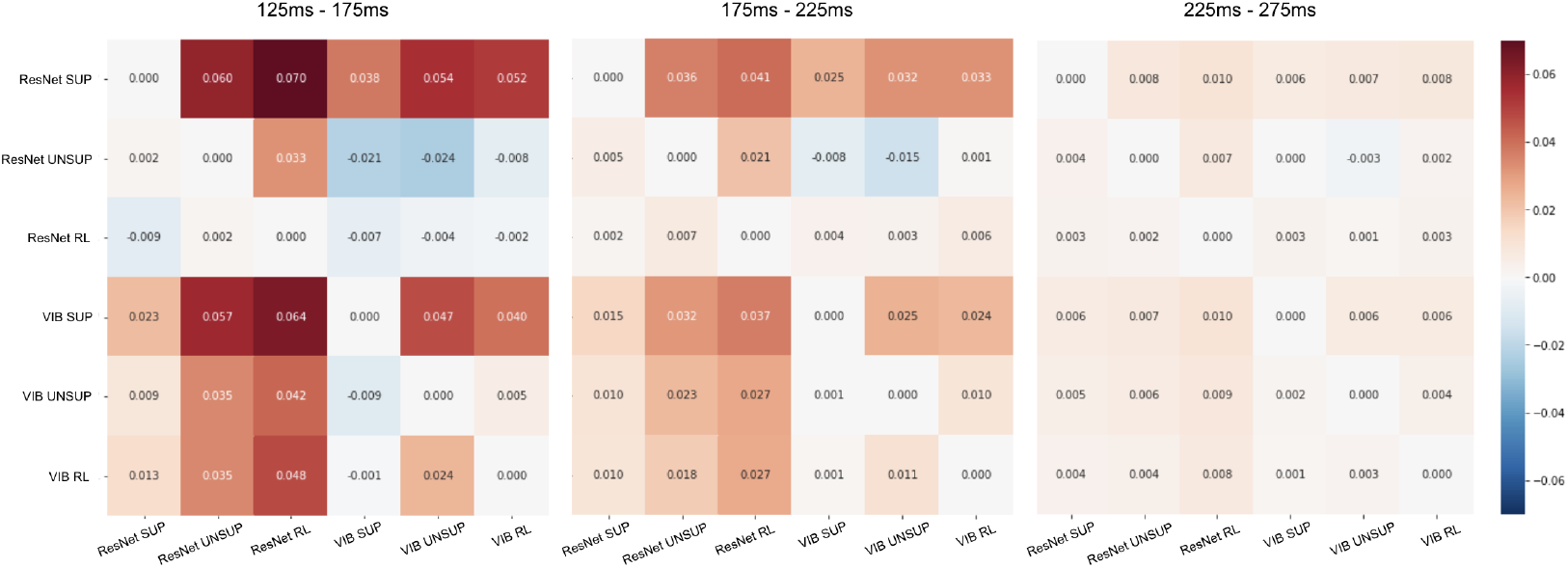
Semi-partial Kendall *τ* correlations. Heatmaps show semi-partial Kendall *τ* between face-selective iEEG RDMs and model RDMs across time windows. Rows denote the predictor (row model’s unique contribution); columns denote the controlled model (partialled out).

**Fig. 5.**
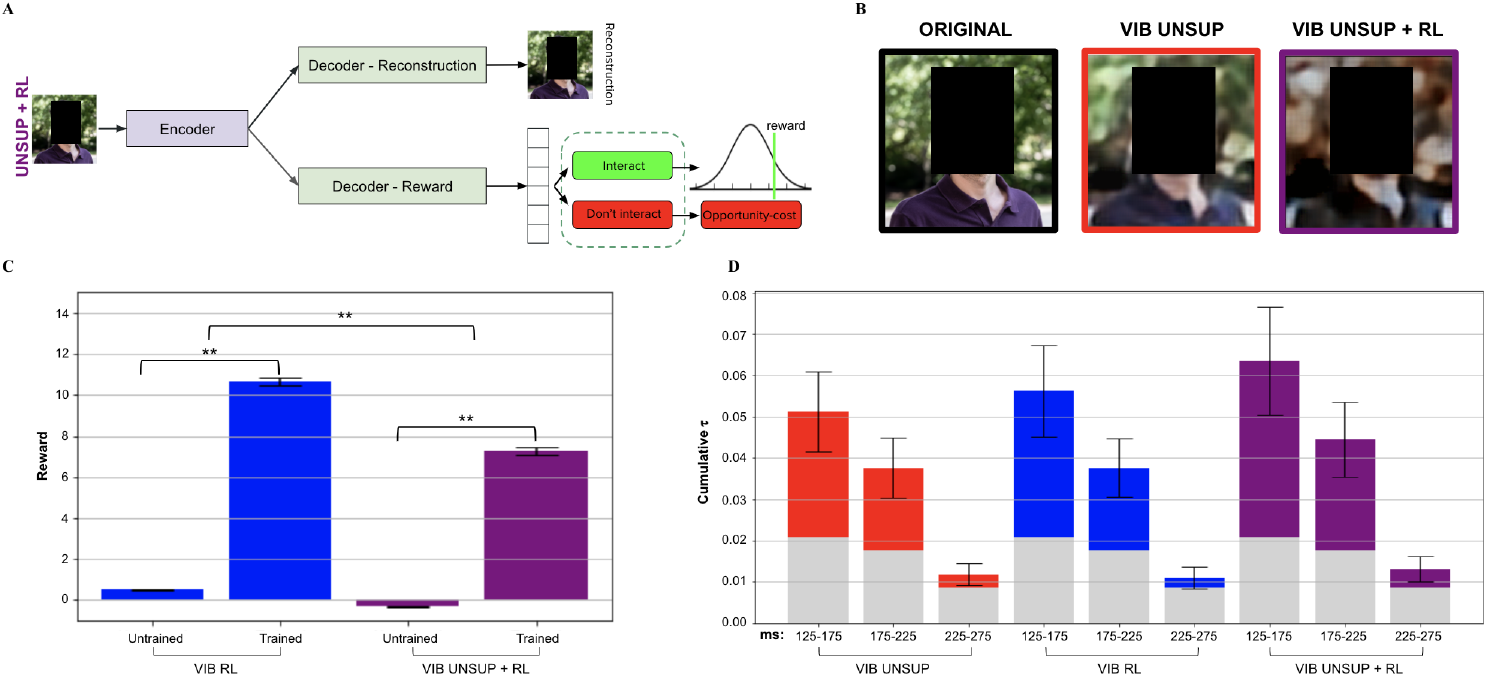
Combination of unsupervised learning and reinforcement learning. **A**. Neural network architectures for the VIB UNSUP+RL model. **B**. Example of image reconstruction from VIB UNSUP model and VIB UNSUP+RL model. **C**. Average reward obtained before and after training for VIB RL model and VIB UNSUP+RL model. **D**. Cumulative Kendall *τ* correlations between face-selective iEEG RDMs and the RDMs from each model averaged over electrodes (*n* = 24).

**Fig. 6.**
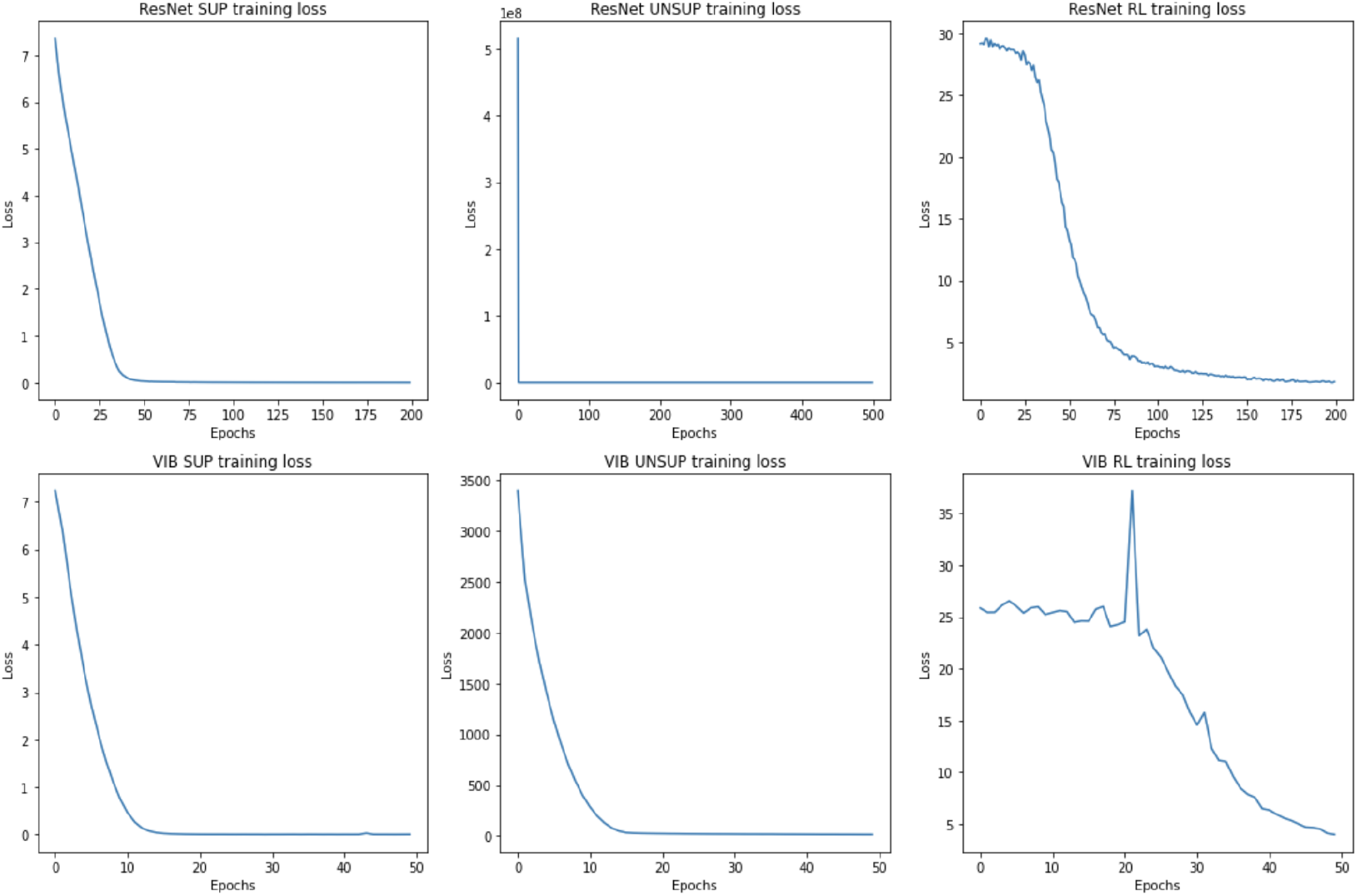
Model training losses.

**Fig. 7.**
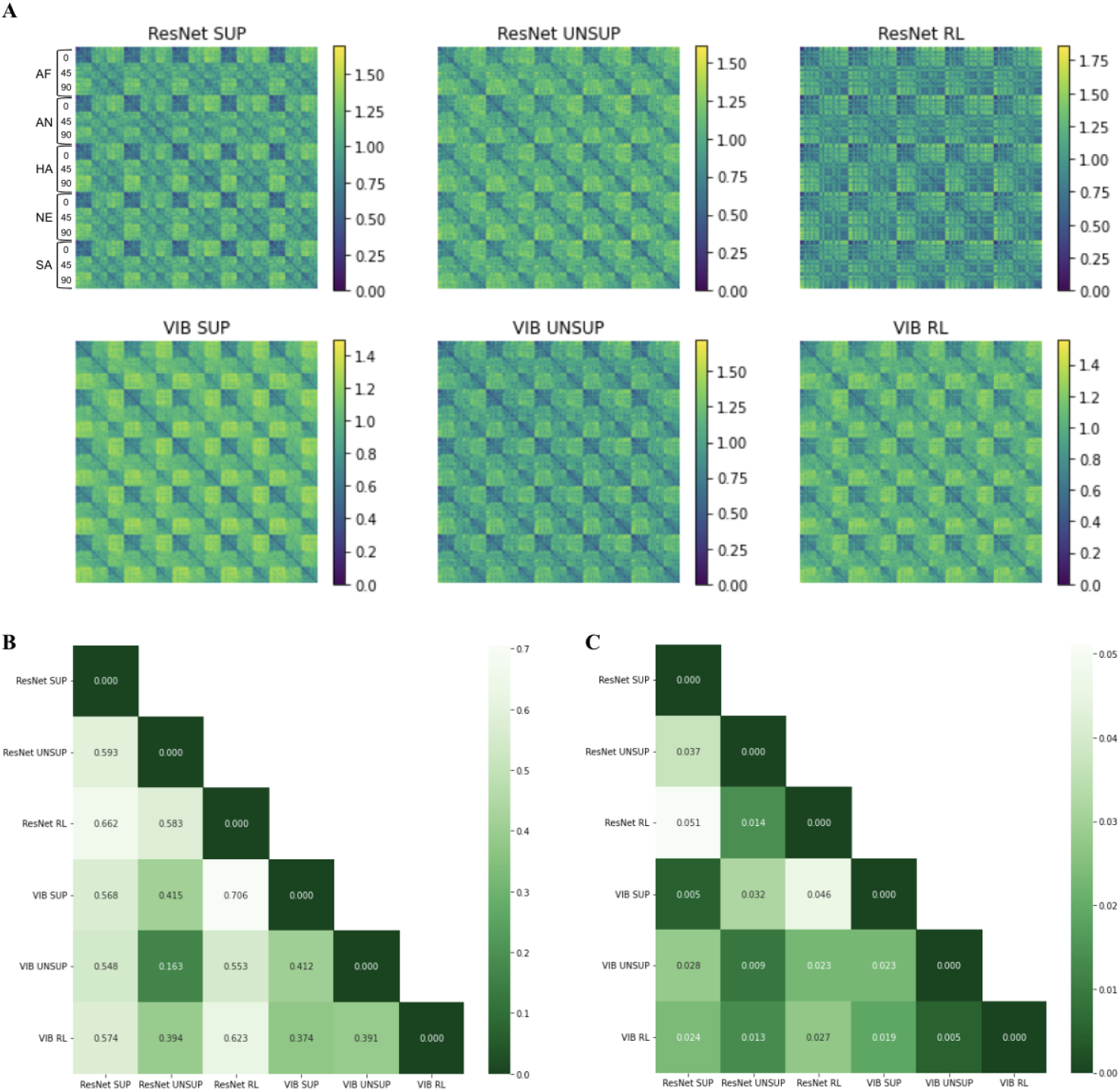
Model comparisons. **A**. RDMs of the latent representations of all models (based on KDEF images used in version B of the experiment). **B**. Kendall’s *τ* ranking distance between the RDMs of the different models. **C**. Euclidean distance between the models’ match with neural responses. The match was computed as cumulative Kendall *τ* values between model RDMs and neural RDMs.

**Fig. 8.**
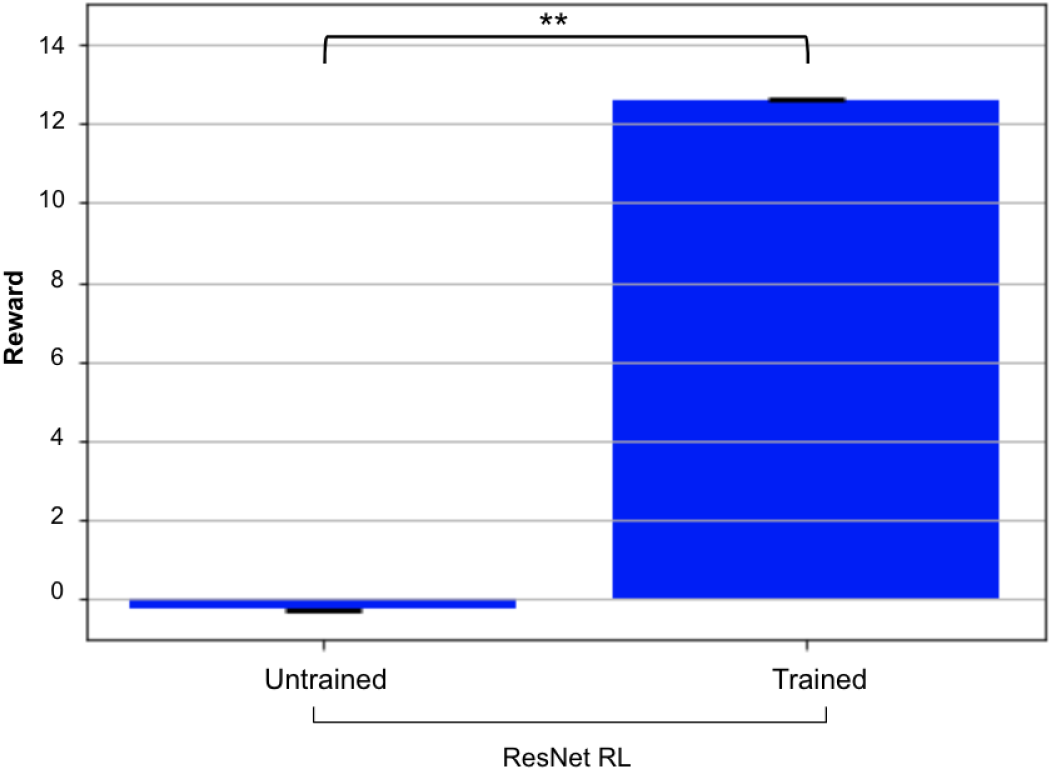
Average reward obtained before and after training for ResNet RL model.

### 3.6 Modeling face representations with a combination of unsupervised learning and feedback from the environment

In realistic settings, rewards are encountered only occasionally. By contrast, unsupervised experience is plentiful. Considering that visual representations are shaped by the feedback an animal receives (Sigala and Logothetis, 2002), but also that unsupervised models have been reported to do well at accounting for neural responses to objects (Zhuang et al., 2021; Konkle and Alvarez, 2022), the visual system might rely on a combination of unsupervised learning and feedback from the environment. Combining these different types of learning, however, can pose challenges. Changes in neural representations that improve performance on the unsupervised task might worsen performance on the approach-avoidance task, and vice versa. To evaluate this, we tested whether a model of face perception that combines these two types of learning can still complete both types of tasks accurately, and whether it acquires representations that correspond more closely to neural responses.

Specifically, we trained a model that uses the VIB DenseNet encoder architecture, but differs from previous models we tested in that the output of the encoder is then fed as input to two decoders (VIB UNSUP+RL, Fig. 5A). The first decoder reconstructs the image (from which we calculated an unsupervised loss), while the second decoder determines whether or not to interact with the person in the image (from which we calculated a reinforcement learning loss). This network is trained to minimize the sum of the two losses, so that the encoder is pushed to learn representations that can be used to reconstruct the original images, and also to perform the approach-avoidance task.

After training this “combined” model, we first aimed to evaluate its performance on the tasks for which it was trained. We compared its ability to reconstruct images to the model trained exclusively to reconstruct images (the unsupervised model), finding that the reconstructions of the combined model were comparatively less accurate, but preserved the overall appearance of the face (Fig. 5B). We also compared the combined model’s ability to maximize reward, finding that the model was able to improve its decisions during training and to obtain significantly larger amounts of reward at the end of the training procedure (Fig. 5C, VIB UNSUP+RL), although the amount of reward it obtained after training was significantly lower than the reward obtained by the model trained exclusively with approach-avoidance (Fig. 5C, VIB RL). In sum, although the combined model did not perform as accurately as the models trained exclusively with one task, it was nonetheless able to perform both tasks.

Next, we evaluated the correspondence between the combined model and neural responses. Compared to the unsupervised model trained exclusively with image reconstruction and to the reinforcement learning model, the combined models’ correlations with neural responses were numerically higher (Fig. 5D). These findings underscore that while the combined model did not perform as accurately as the models trained exclusively with one task, it nonetheless showed a performance that was at least equivalent to that of the VIB UNSUP and VIB RL models in accounting for neural responses in face-selective electrodes, with a non-significant main effect for the model type (*F* (2, 213) = 0.484, *p* = 0.617). More evidence will be needed to determine whether combining unsupervised learning and feedback from the environment can yield representation that provide a closer match to neural responses.

## 4 Discussion

Deep network models trained based on feedback from the environment (RL) were able to account for neural responses comparably for both supervised and unsupervised models. This is particularly significant considering that the supervised models tested in this study were trained to recognize face identity: such supervised models have been shown to perform well at capturing face-selective neural responses, outperforming supervised models trained with other tasks such as facial expression classification (Schwartz et al., 2023a).

Unlike SUP models, that require ground-truth labels which are not usually available in realistic settings, RL models can learn through more realistic interactions with the environment. In addition, unlike UNSUP models, the representations in RL models do not depend exclusively on the input stimuli, but also on the reward the models receives from the environment. In this respect, VIB RL models can account for previous results showing that the visual system is shaped by animals’ interactions with the environment and by the tasks they are trained to perform (Sigala and Logothetis, 2002; De Baene et al., 2008).

Models trained based on feedback from the environment were only able to obtain comparable performance to supervised and unsupervised models when using a novel encoder architecture, that combines the strengths of variational encoders (Kingma and Welling, 2013) and of densely connected neural networks – DenseNets (Huang et al., 2017). By contrast, when using a ResNet architecture (He et al., 2016), representations learned with a supervised identity classification task correlated more strongly with neural responses compared to the representations learned with the other models tested (UNSUP and RL).

Recent work found that deep RL models such as deep Q networks can capture neural responses across multiple brain regions while participants play Atari games (Cross et al., 2021). In addition to correlating with neural responses in frontal and parietal regions, the models’ representations also correlated with responses in regions in ventral temporal cortex known to encode perceptual representations (Cross et al., 2021). Temporal and frontal regions interact during reinforcement learning (Gershman and Daw, 2017). Previous studies focused on Atari games that display simple stylized visual stimuli. Therefore, a model could capture neural responses to such stimuli, but might still be unable to account for responses to more complex and nuanced stimuli like human faces. While the present work uses a different model architecture (VIB DenseNet), it indicates that models that learn based on feedback from the environment have the potential to account for the representation of naturalistic images.

The task performed by the RL model, while being more realistic than categorization based on ground-truth labels, is still a simple approximation of real-life interactions. Despite this, the VIB RL model was still able to account for neural responses comparably to the VIB SUP and VIB UNSUP models. This finding suggests that, when using the VIB DenseNet encoder, RL models are a viable approach to capture perceptual representations. We hypothesize that improving the realism and complexity of the reinforcement learning task in future studies will also improve the correspondence between the models’ representations and neural responses.

The architecture of the models had an impact on the results. Having been trained on the same dataset, models employing the ResNet encoder architecture and models employing the VIB DenseNet encoder architecture demonstrated a different degree of correspondence with neural responses. This was especially true in the RL models: when training on the same dataset and task, the VIB RL model showed significantly higher correlations with neural responses than the ResNet RL model. Two aspects of the architecture differed between the two models. First, unlike the ResNet RL model, the VIB RL model is based on a combination of DenseNet and variational autoencoders (VAEs) architecture. Second, the VIB RL model includes a probabilistic bottleneck, which injects stochasticity during training, and a regularization term in the loss (the KL-divergence). Either of these features (or both) could have contributed to the difference in performance. The task the networks were trained on (and the associated decoder architecture) also had an impact on the results, even when the encoder architecture was matched. In fact, both the ResNet UNSUP model and the ResNet RL model did not perform as well as the ResNet SUP model, showing that the training task (and the associated decoder) contributed to the correspondence between models and neural responses even when the encoder architecture was held constant.

We anticipated the VIB UNSUP+RL combined model to exhibit greater correspondence to neural responses, but its correlation with neural data was not significantly different from that of the VIB UNSUP model and the VIB RL model. The combined model’s reconstruction performance and the reward it achieved after training were lower compared to the models trained specifically for one single task. Within the multi-task model, the optimization process for one task might have affected negatively the optimization of the other task (Sener and Koltun, 2018). This finding encourages the search for other approaches to combine unsupervised learning and reinforcement learning that can perform well at both tasks. Such approaches might also yield representations that provide a closer match to neural responses.

Our analysis also revealed insights about the differences between neural representations in ventral and lateral temporal regions. Thanks to the temporal resolution of iEEG, we were able to investigate differences between the two streams across multiple time windows. First, we found a significant variation in the correspondence of these regions with the models across different time windows. Second, we observed that overall the models’ correspondence with ventral electrodes was greater than with lateral ones, particularly in the initial time windows. This disparity might be due to the nature of the stimuli used to train our DCNNs. Previous research has reported that lateral regions have stronger responses to dynamic stimuli (Pitcher et al., 2019), whereas our models were trained on static images. This might account for the lower correspondence observed with the lateral electrodes. This discrepancy could potentially be addressed by making use of deep networks that can process dynamic stimuli (Lotter et al., 2016; Feichtenhofer et al., 2019).

## Notes

https://github.com/sccnlab/facefeedback/tree/main

### Competing Interest Statement

The authors have declared no competing interest.

